# Towards the prediction of non-peptidic epitopes

**DOI:** 10.1101/2021.06.09.447727

**Authors:** P. F. Zierep, R. Vita, N. Blazeska, J. A. Greenbaum, B. Peters, S. Günther

## Abstract

In-silico methods for the prediction of epitopes can support and improve workflows for vaccine design, antibody production, and disease therapy. So far, the scope of B cell and T cell epitope prediction has been directed exclusively towards peptidic antigens. Nevertheless, various non-peptidic molecular classes can be recognized by immune cells. These compounds have not been systematically studied yet, and prediction approaches are lacking. The ability to predict the epitope activity of non-peptidic compounds could have vast implications; for example, for immunogenic risk assessment of the vast number of drugs and other xenobiotics. Here we present the first general attempt to predict the epitope activity of non-peptidic compounds using the Immune Epitope Database (IEDB) as a source for positive samples. The molecules stored in the Chemical Entities of Biological Interest (ChEBI) database were chosen as background samples. The molecules were clustered into eight homogeneous molecular groups, and classifiers were built for each cluster with the aim of separating the epitopes from the background. Different molecular feature encoding schemes and machine learning models were compared against each other. For those models where a high performance could be achieved based on simple decision rules, the molecular features were then further investigated. Additionally, the findings were used to build a web server that allows for the immunogenic investigation of non-peptidic molecules (http://tools-staging.iedb.org/np_epitope_predictor). The prediction quality was tested with samples from independent evaluation datasets, and the implemented method received noteworthy Receiver Operating Characteristic-Area Under Curve (ROC-AUC) values, ranging from 0.69-0.96 depending on the molecule cluster.

## Introduction

Defense against pathogens via the adaptive immune system depends on the distinction between endogenous and exogenous molecules produced by the host and pathogen, respectively. This distinction is made by receptors located on the surface of T and B lymphocytes. The specific part of an antigen that interacts with the T cell receptor (TCR) or B cell receptor (BCR) is known as the epitope.

T cells recognize antigens bound to the major histocompatibility complex (MHC) presented on the surface of cells. All nucleated cells present endogenous antigens via MHC class I molecules as a self/non-self distinction feature. Professional antigen-presenting cells, such as macrophages and B cells, present antigens primarily derived from the extracellular space via MHC class II molecules. B cell recognition is mediated by receptors located on the cell membrane. Activated B cells differentiate into plasma cells, which can secrete a soluble form of their receptors as antibodies. Antibodies can impede the function of pathogens or tag the pathogen for elimination by macrophages. Specific antibodies with targeted recognition are widely used for the development of vaccines [1], therapeutic antibodies [2], immunodiagnostic tools [3], and immunoassays [4–6].

The vast majority of known epitopes are derived from proteins. However, peptides are not the only entities that can be detected by the immune system. In fact, there are other molecular classes that elicit an immune response, such as lipids, carbohydrates, drugs, and metals [7]. Small molecular entities, such as metals (e.g., nickel) and organic compounds (e.g., aniline and its derivatives) are referred to as haptens. They must conjugate with larger carrier proteins to be recognized by T cells or specific antibodies. Larger molecular entities, such as polysaccharides [8,9], glycolipids [10], and lipids [10,11], can lead to an immune response directly. Cross-reactive carbohydrate determinants play a major role in allergic disease and anaphylactic events [12,13].

Recognition of epitopes associated with pathogens (e.g., bacteria, virus, fungi) leads to the protection of the host from further exposure. However, unwanted immunogenicity can lead to serious health problems for the host. When natural or synthetic compounds, derived from food, cosmetics, or plants, are recognized by the immune system, an allergy may occur, which can lead to symptoms such as skin inflammation and asthma. The immune response against therapeutics, mediated by so-called anti-drug antibodies, can decrease or even reverse the effects of the drug [14]. Furthermore, induced autoimmune responses can also be directed against the body’s own biomolecules. Although more than 100 different autoimmune diseases are described [15], their exact causes are mostly unknown.

The prediction of possible epitopes is a crucial step towards the development of novel vaccines and drugs, prevention of allergic reactions, and can also play an important role in the general understanding of the immune system. Various approaches for the prediction of peptidic epitopes have been described [16,17]. Most of these prediction approaches use known peptidic epitopes to generate rule-based or machine learning-based classifiers. Consequently, the bottleneck for efficient epitope prediction is created by the availability and quality of known epitopes. The Immune Epitope Database (IEDB) is a continuously updated large collection of literature-derived epitopes [18], which has been the source of training samples for various peptide-based epitope prediction tools.

To the best of our knowledge, the immunogenic recognition of non-peptidic compounds has not yet been studied systematically and, thus far, no method has been described to predict non-peptidic epitopes. The investigation of statistically significant characteristics of non-peptidic epitopes could have vast implications for the development of novel materials, cosmetics, and drugs. The in-silico prediction of epitope activity of non-peptidic molecules would allow for risk assessment prior to labor-intensive experimental assays. An immunogenicity prediction tool could help to streamline drug development pipelines and synthetic material evaluation progress, in a similar way to toxicity prediction tools [19]. Furthermore, various allergic and autoimmune reactions may be associated with xenobiotics present in the environment [20]. Their immunogenic reevaluation could help to locate the source of related, globally increasing health problems [21].

The largest collection of curated non-peptidic epitopes exists in the IEDB, where more than 2700 non-peptidic structures with reported positive B cell and/or T cell assays are described. These molecules were used as positive samples and compared against background molecules from the Chemical Entities of Biological Interest (ChEBI) database [22]. Different molecular encoding schemes and machine learning models were benchmarked for their ability to predict the epitope activity of non-peptidic molecules. The findings were compiled into a prediction web server, which allows for the thorough immunogenic assessment of non-peptidic molecules.

## Results

### Clustering into homogeneous molecular subsets

The molecules stored in the IEDB (2,719 positive epitope samples) were merged with the ChEBI molecules (42,643 background samples) and converted into bit features using the Morgan fingerprints algorithm [23], and clustered into homogenous molecular subsets using k-means clustering. The total number of clusters was determined by plotting of the cluster inertia (i.e., density of the clusters) against the number of clusters. The appearance of a kink in the plot (elbow method) would indicate an ideal cluster number (see Figure 1).

**Figure 1.**
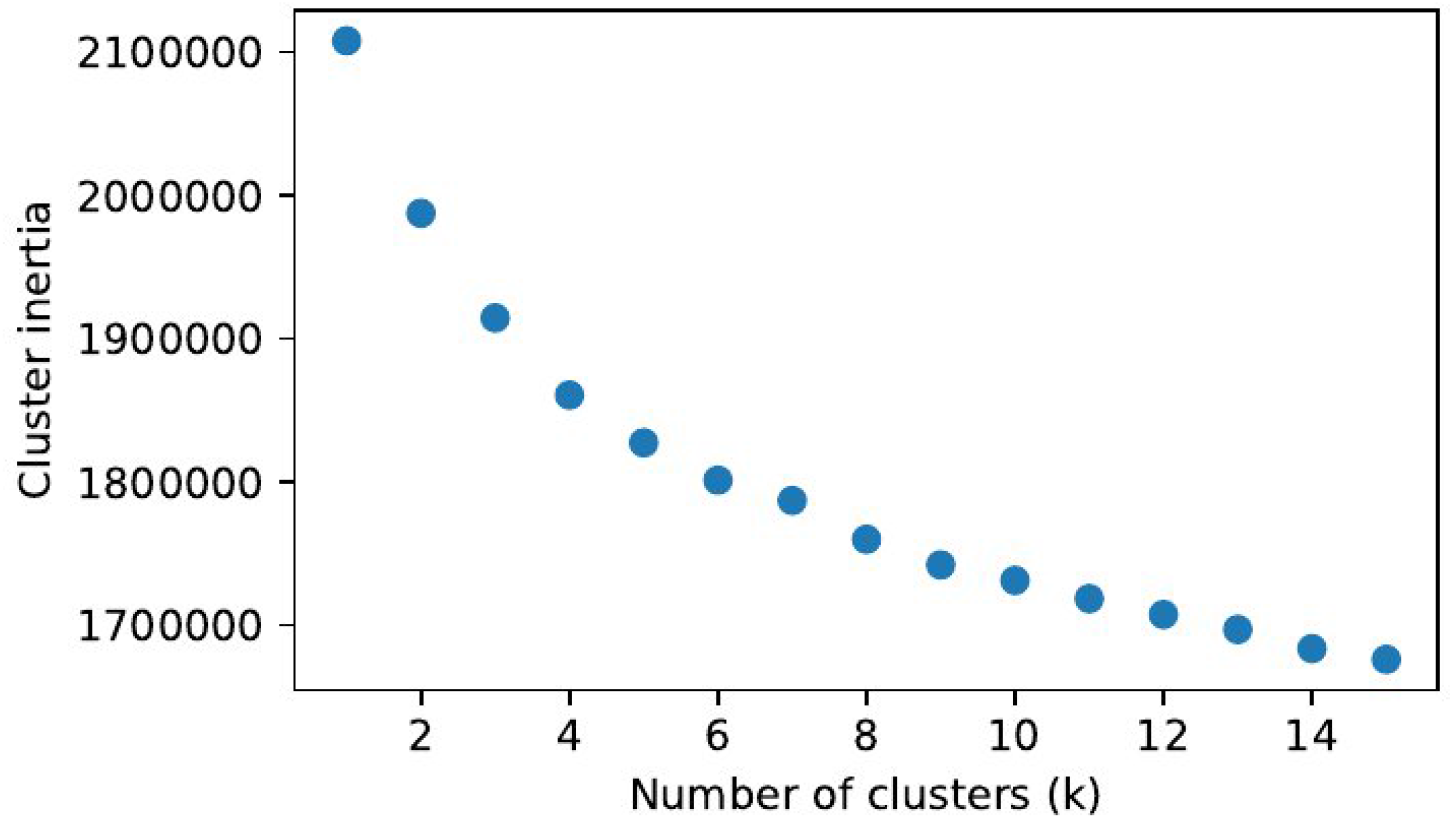
Cluster inertia plotted against the number of clusters (k). The cluster inertia is computed as the sum of squared distances of samples to their closest cluster.

Even though an unambiguous kink is not observed, the cluster inertia decreases only marginally for more than 8 clusters. This cluster number was chosen and the individual clusters were further described. The principal component visualization of the clusters (Figure 2) shows that the clusters overlap and have different sizes.

**Figure 2.**
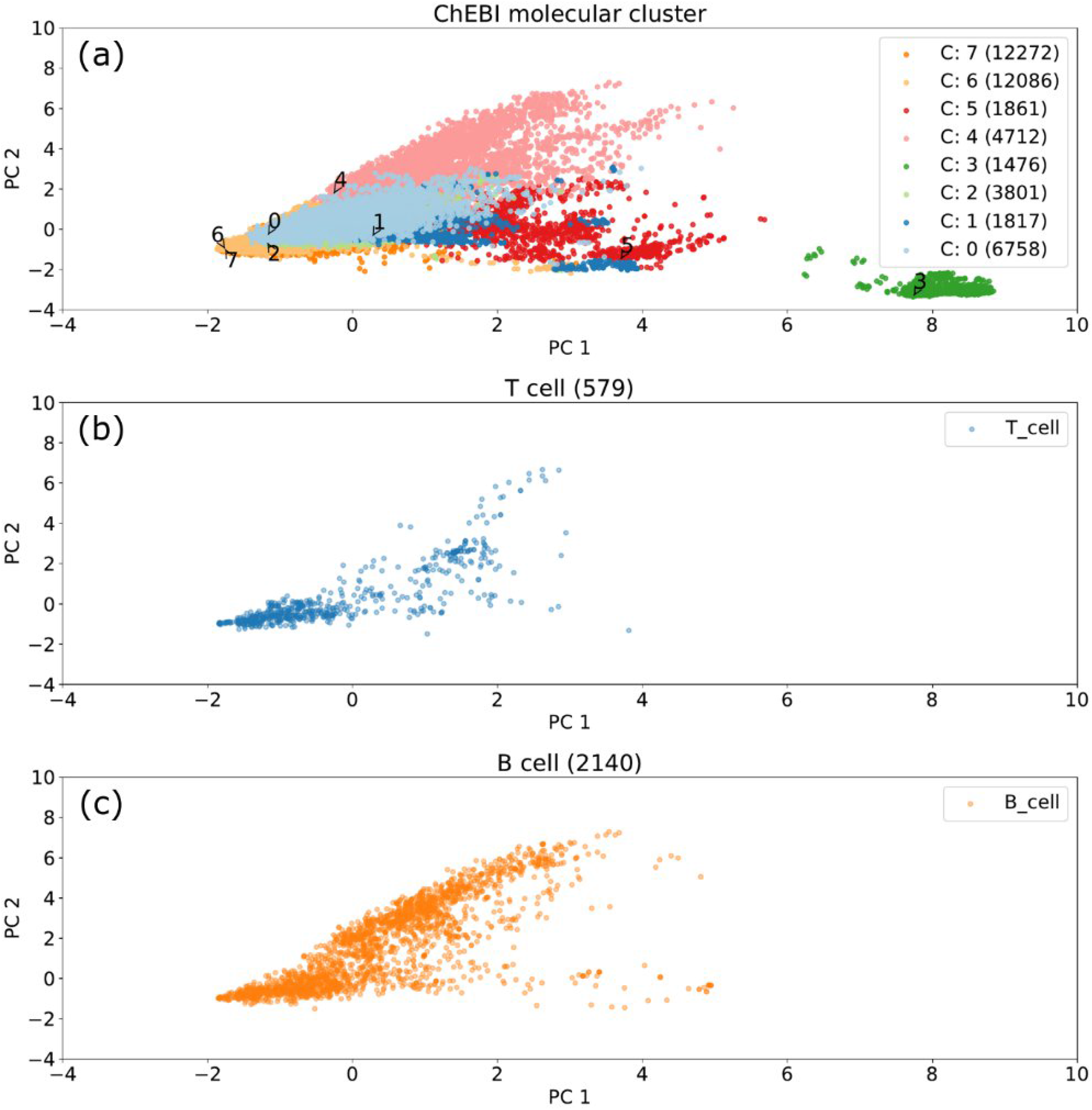
Principal component visualization of the ChEBI dataset. (a) Principal components of the 8 clusters and their sizes. (b) T cell epitopes. (c) B cell epitopes.

Epitopes that tested positive in B and T cell assays are present in all clusters except for Cluster 3, which contains only non-epitopes. Cluster 3 comprises exclusively Coenzyme A (CoA)-derived molecules. Example molecules for each cluster are shown in Figure 3. The clusters were described by an ontology enrichment analysis as output of the BiNChE (acronym not described in original publication) web tool [24]. See Methods for a detailed explanation of the BiNChE tool and output. An example of the analysis for cluster 4 is shown in Table 1. For this cluster, all enriched ontology terms were related to glucoside and oligosaccharide molecules; therefore, the cluster name “glucoside/oligosaccharide derivatives” was chosen. Similar cluster names were derived for all clusters shown in Table 2. The BiNChE analysis for all clusters is shown in the Supplementary Data.

**Table 1.** BiNChE ontology analysis of cluster 4. The name “glucoside/oligosaccharide derivatives” was chosen for this cluster.

| ChEBI ID | ChEBI Name | Fold-enrichment | Sample coverage (%) |
| --- | --- | --- | --- |
| CHEBI:22485 | glucosamine oligosaccharide | 23.71 | 10 |
| CHEBI:22483 | amino oligosaccharide | 22.51 | 18 |
| CHEBI:63563 | oligosaccharide derivative | 21.27 | 27 |
| CHEBI:63353 | disaccharide derivative | 19.46 | 9 |
| CHEBI:22798 | beta-D-glucoside | 18.45 | 11 |
| CHEBI:35436 | D-glucoside | 18.32 | 12 |
| CHEBI:60980 | beta-glucoside | 18.24 | 11 |
| CHEBI:24278 | glucoside | 17.93 | 12 |
| CHEBI:35313 | hexoside | 17.30 | 13 |
| CHEBI:33563 | glycolipid | 12.97 | 17 |

**Table 2.** Summary of the compiled molecular clusters. The mean fold-enrichment can be used as an indicator of the homogeneity of the cluster.

| Cluster | Name | Mean Fold-enrichment | Number of molecules | B cell | T cell | Fingerprint size |
| --- | --- | --- | --- | --- | --- | --- |
| 0 | steroid/terpenoid like | 9.33 | 6,758 | 191 | 57 | 5,942 |
| 1 | betaine/glycerolipid derivatives | 17.28 | 1,817 | 36 | 31 | 634 |
| 2 | fatty acid derivatives | 20.59 | 3,801 | 59 | 56 | 1,381 |
| 3 | acyl-CoA derivatives | 69.73 | 1,476 | 0 | 0 | 606 |
| 4 | glucoside/oligosaccharide derivatives | 19.01 | 4,712 | 1,079 | 106 | 3,636 |
| 5 | nucleobase-containing molecular entities | 9.64 | 1,861 | 101 | 15 | 1,167 |
| 6 | diverse small molecules | 2.49 | 12,086 | 252 | 94 | 2,267 |
| 7 | cyclic halide/phenols | 3.37 | 12,272 | 422 | 220 | 5,370 |

**Figure 3.**
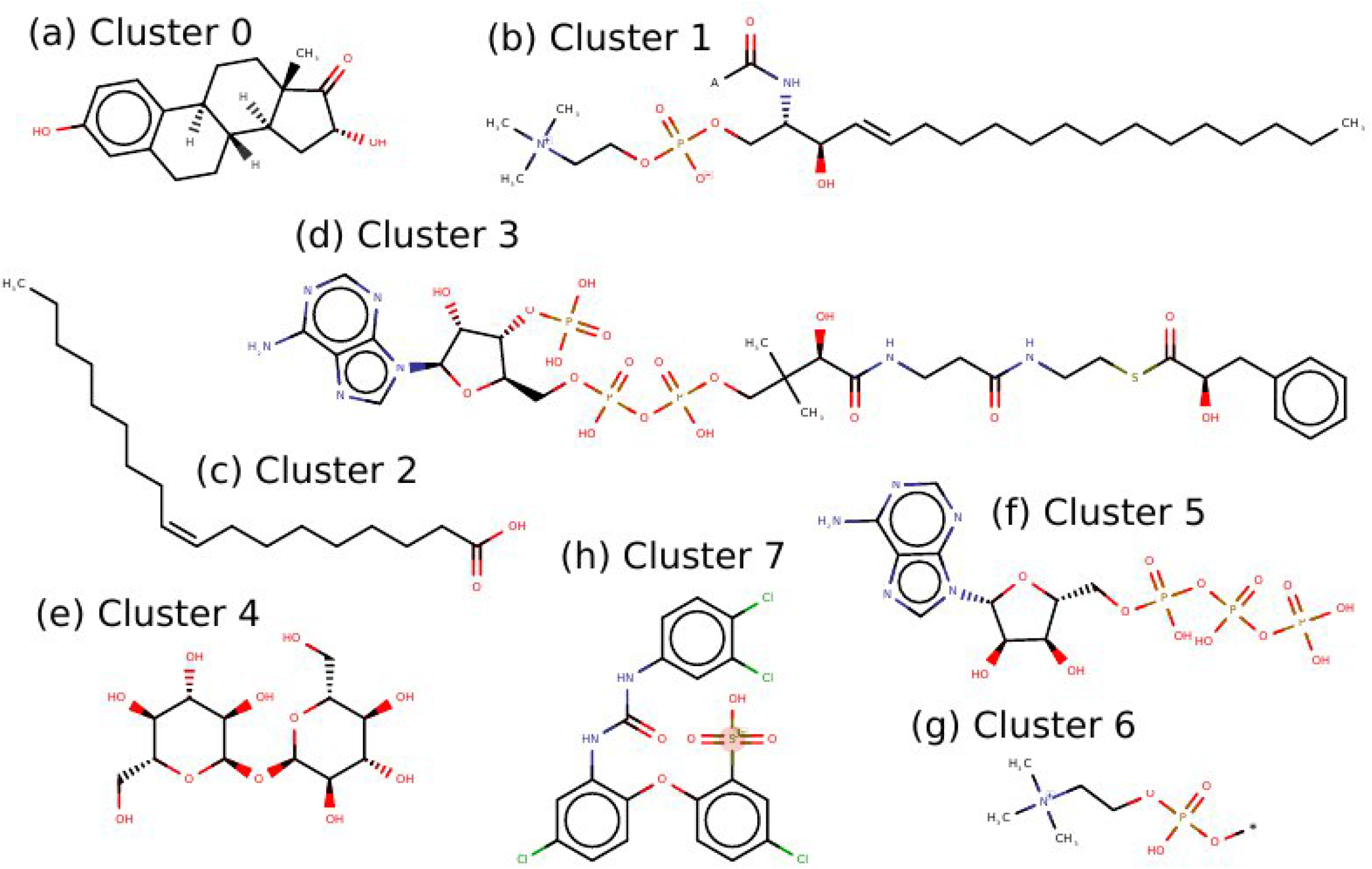
Example molecules for each cluster generated for the ChEBI dataset. ChEBI IDs used for the example molecules: (a) Steroid/terpenoid like: CHEBI:776; (b) Betaine/glycerolipid derivatives: CHEBI:17636; (c) Fatty acid derivatives: CHEBI:16196; (d) Acyl-CoA derivatives: CHEBI:11010; (e) Glucoside/oligosaccharide derivatives: CHEBI:16551; (f) Nucleobase-containing molecular entities: CHEBI:15422; (g) Diverse small molecules: CHEBI:55395; (h) Cyclic Halide / Phenols: CHEBI:59246. All examples represent molecules that have been tested positive in B cell essays - except for the acyl-CoA derivatives, where no epitope was described.

An intuitive metric to describe the cluster homogeneity is given by the fold-enrichment (ratio between the enrichment in the selected samples and enrichment in the background samples) term of the BiNChE analysis. Clusters that can be separated from the main dataset show very high fold-changes of their associated ontology terms. The mean fold-enrichment of the 10 most significant ontology terms are added to Table 2. Most clusters can be distinguished, since their ontology terms are highly enriched - except for clusters 6 and 7 which do not allow a clear cluster description.

The diversity of the clusters is represented by the number of automatically generated Morgan fingerprint features and is shown in Table 2. Cluster 7 generated the most features with an array size of 63 × 10^6^ values (11,805 samples x 5,370 features). The entire ChEBI database would require 615 × 10^6^ values (42,643 samples x 14,583 features).

### Performance of different Morgan fingerprint parameters

The radius and chirality options for the Morgan fingerprint generation were analyzed with regard to the epitope prediction performance of random forest (RF) classifiers. Random forest classifiers represent a class of machine learning models, that are widely used for novel classification tasks due to their general applicability. The performance was evaluated using the Receiver Operating Characteristic Area Under Curve (ROC-AUC) metric. See Methods for a detailed explanation of the machine learning models and performance metrics. The classification performance was slightly better using chiral fingerprints as compared to non-chiral fingerprints (ROC-AUC difference of 0.01 – 0.02) for all radii parameters.

The performance using the chiral fingerprints for each radii parameter is shown in Figure 4. All clusters show the poorest performance when the fingerprints are generated with the radius option 0. The substructures generated with this option only include atom type and connectivity information. The performance increases in many cases with higher radii, although this tendency cannot be observed for all clusters. Cluster 1 and 2 show the strongest fluctuation regarding the radii parameter.

**Figure 4.**
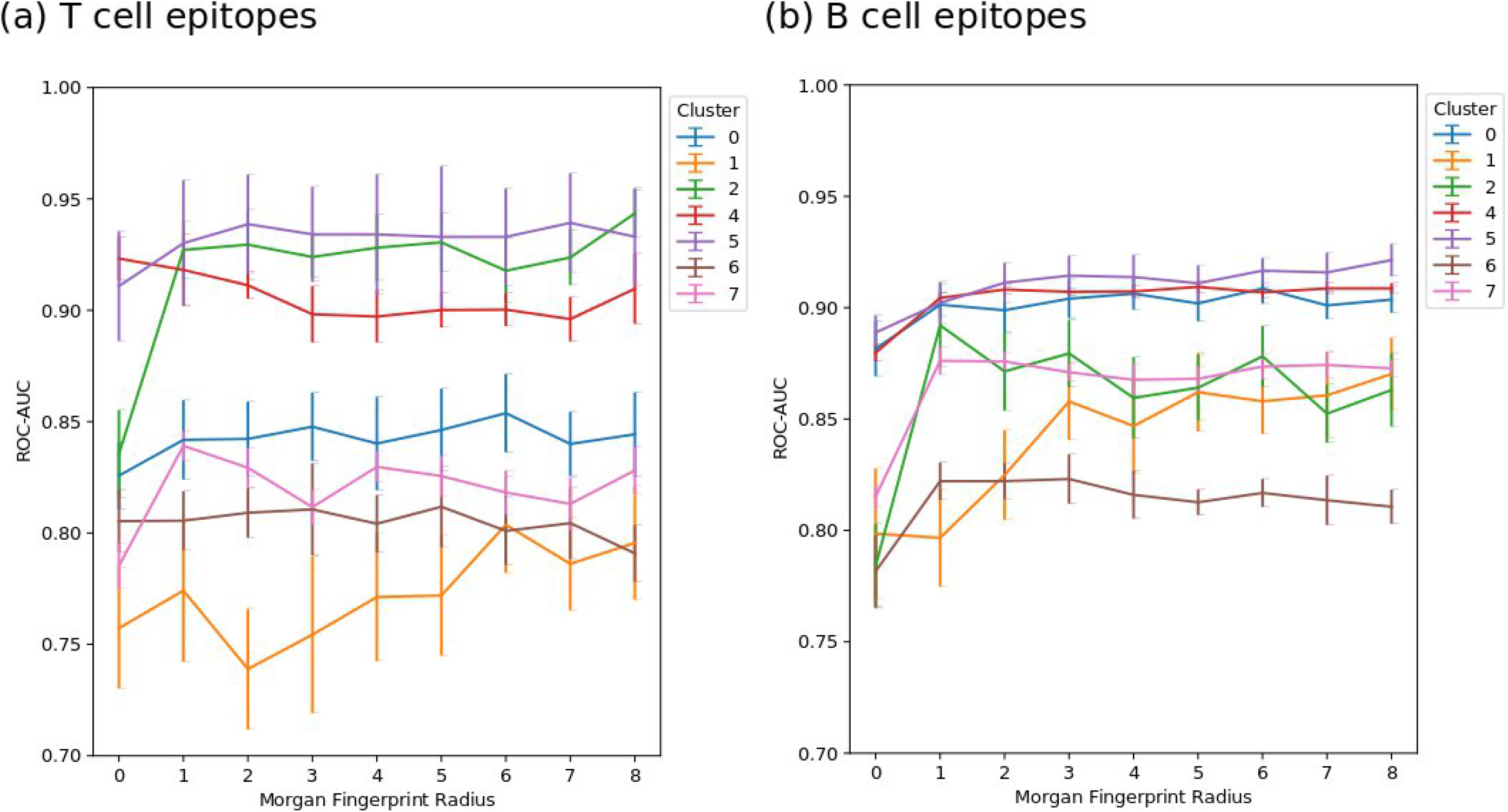
Cross-validation performance of the RF models for different radii parameters used to generate Morgan fingerprints. The prediction of epitopes that tested positive in T cell assays (a) and B cell assays (b).

Chiral fingerprints with a radius of 3 were chosen for the following model benchmark and comparison with Tanimoto similarity-based reference classifiers.

### Model performance

The epitope prediction performance of different machine learning models were compared. The model performance for the prediction of B cell epitopes is shown in Figure 5 and the prediction of T cell epitopes in Figure 6. The RF and neural network (NN) models performed similarly for most molecular clusters. Both models outperformed the k-nearest neighbor (k-NN) models. RF models, trained on randomly assigned positive samples (referred to as dummy models), yielded an ROC-AUC close to 0.5 for all feature sets.

**Figure 5.**
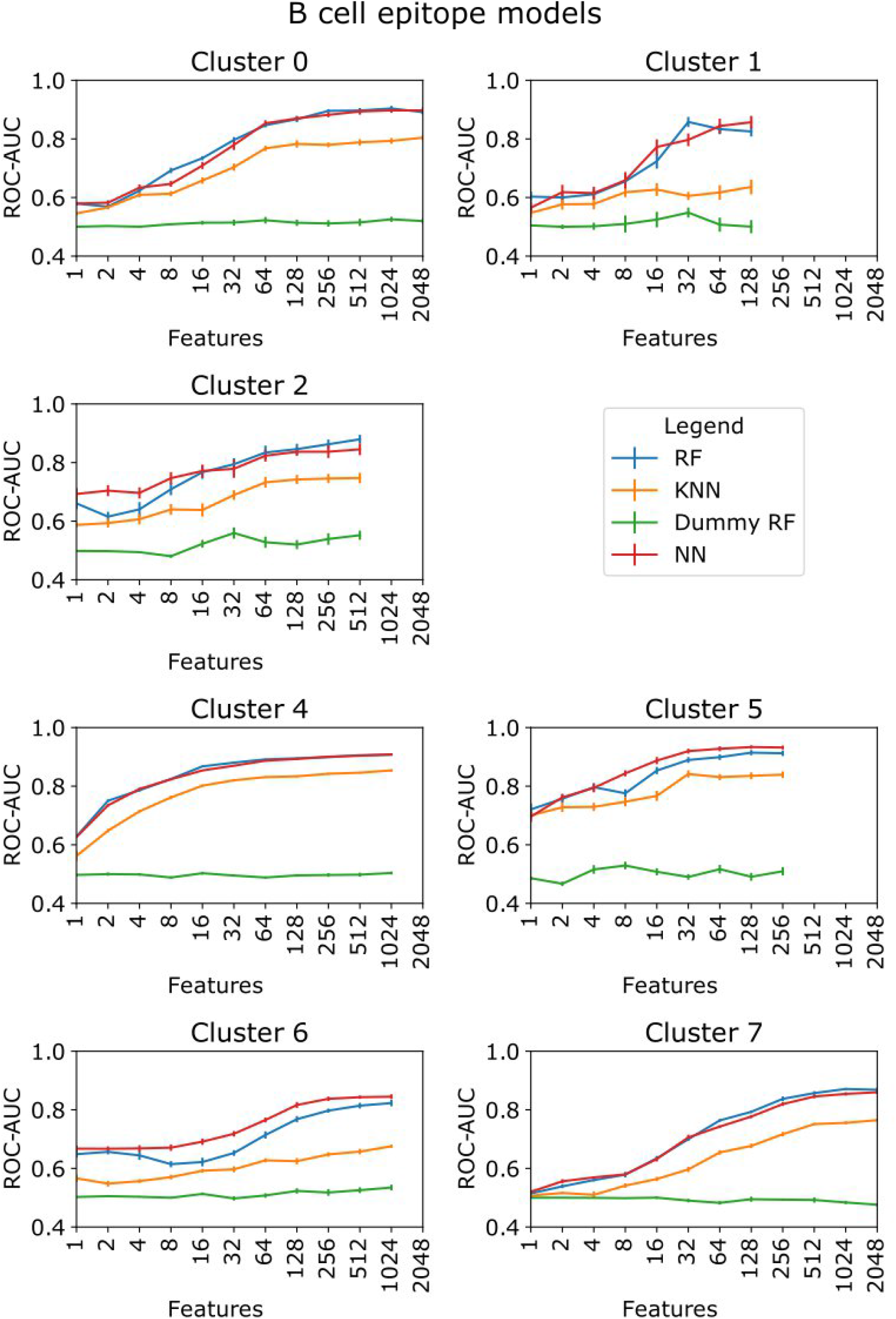
Model comparison for different feature sets for the epitopes that tested positive in B cell assays. Cluster 3 was not benchmarked, since there were no epitopes known for this cluster.

**Figure 6.**
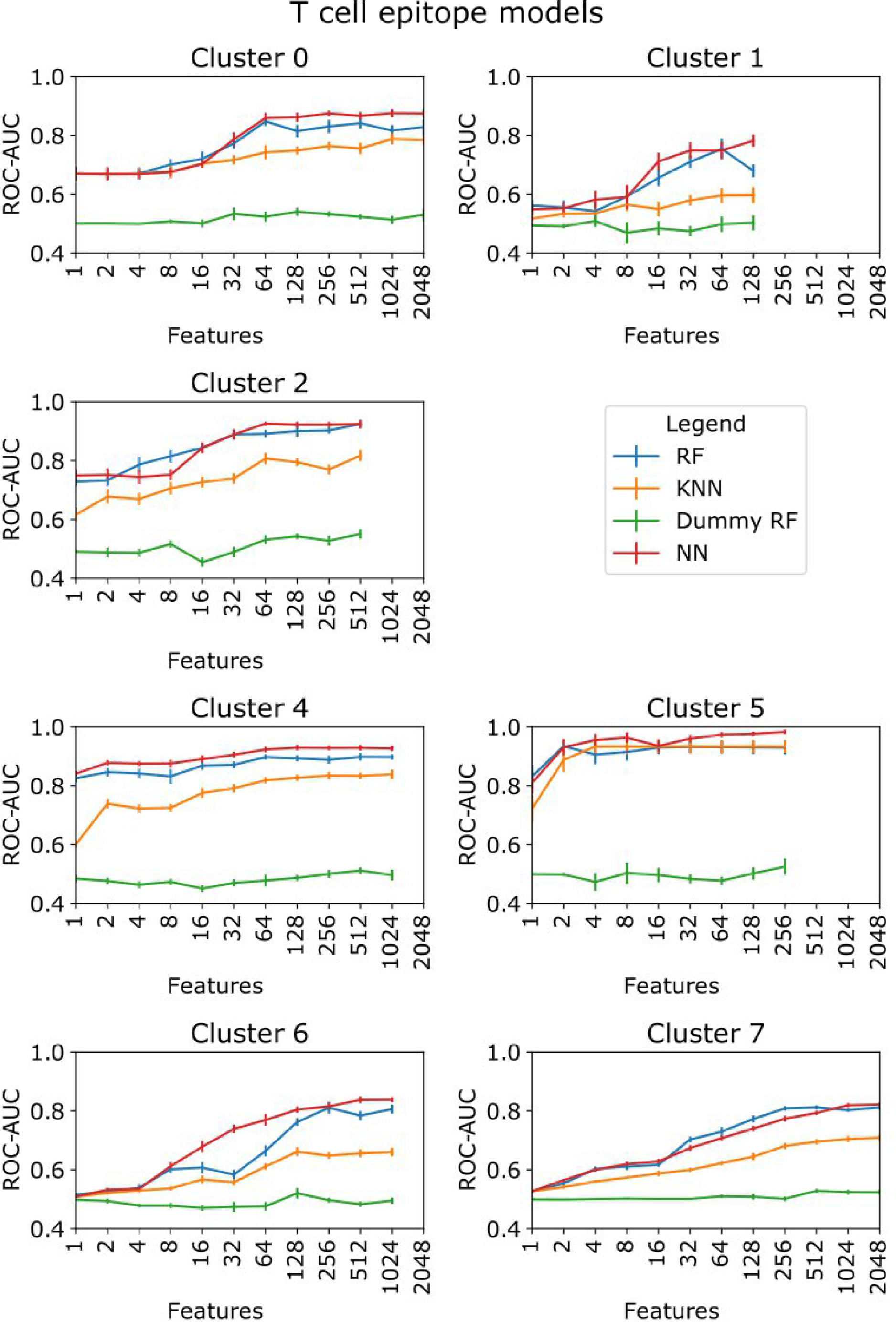
Model comparison for different feature sets for the epitopes that tested positive in T cell assays. Cluster 3 was not benchmarked, since there were no epitopes known for this cluster.

### Epitope prediction

The RF classifiers were compared to Tanimoto similarity-based classifiers. The cross-validation benchmark for each cluster and assay type is shown in Figure 7. It can be observed that, given enough features, the RF models can separate epitopes from the background molecules with high ROC-AUC scores of at least 0.8 for all clusters. In all cases, the RF models yield at least similar ROC-AUC values or outperform the similarity models.

**Figure 7.**
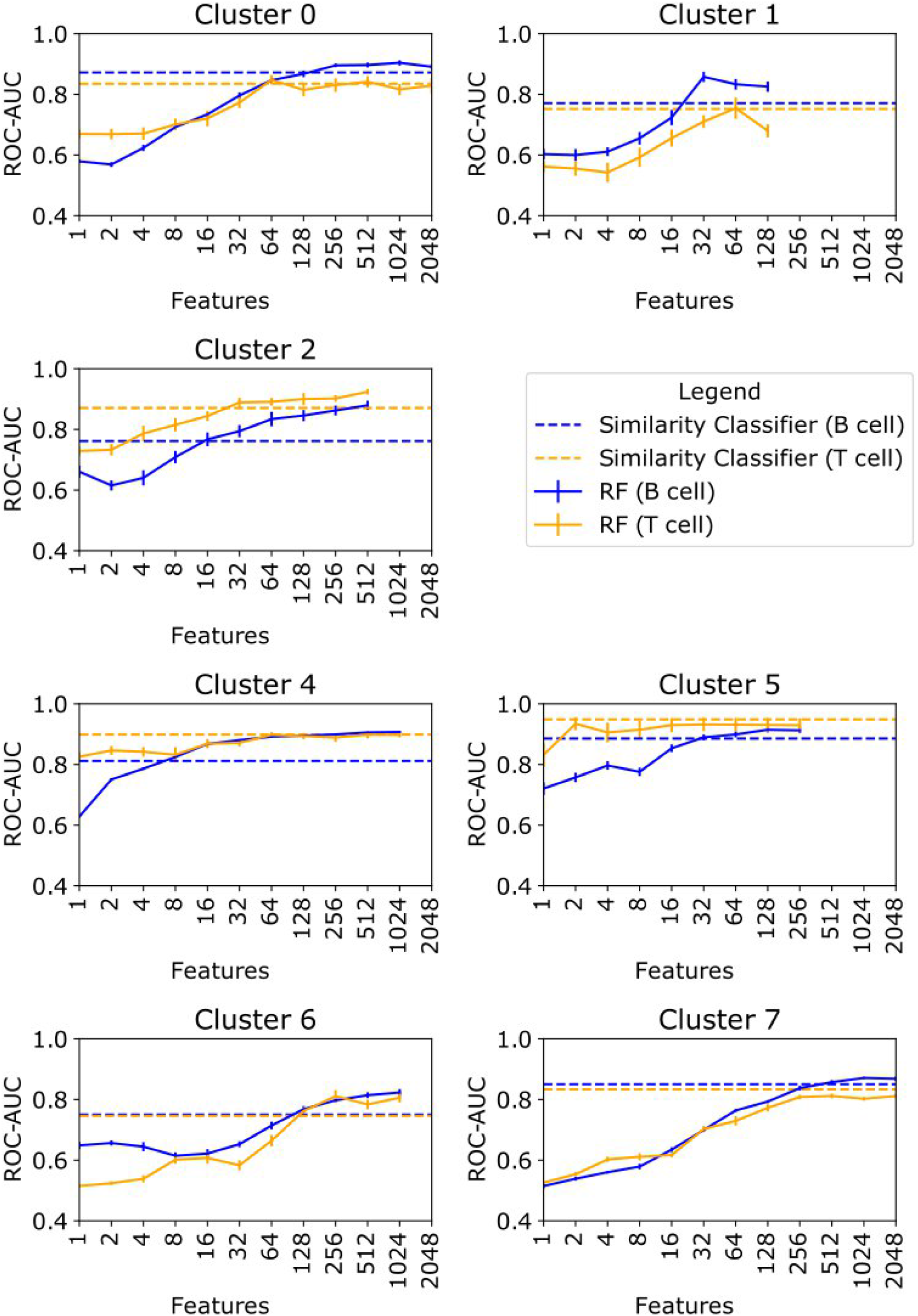
Performance of the epitope classifiers for different feature sets. Cluster 3 is not benchmarked, since there are no epitopes known for this cluster. The RF classifiers are depicted with a continuous line and the similarity classifiers are shown with a dotted line.

Most remarkable are those RF models that yielded high ROC-AUC (> 0.8) values even with low feature sets (clusters 4 and 5 and the T cell epitopes of cluster 2). For these models, the prediction logic can be interpreted based on a few features. The related substructures were investigated in detail in the following section.

The best RF models were used to predict the samples from the independent test dataset of molecules that were not used for the initial training of the classifiers. The performance on the test dataset is shown in Table 3.

**Table 3.** Epitope prediction performance of the RF models on the test dataset. The ROC-AUC values could not be computed for clusters without positive samples. The ROC-AUC value for all clusters takes every prediction into account.

| Cluster | # B cell epitopes | B cell ROC-AUC | # T cell epitopes | T cell ROC-AUC | ChEBI Background |
| --- | --- | --- | --- | --- | --- |
| 0 | 3 | 0.76 | 6 | 0.72 | 255 |
| 1 | 0 | - | 0 | - | 5 |
| 2 | 0 | - | 11 | 0.78 | 113 |
| 4 | 31 | 0.69 | 24 | 0.91 | 894 |
| 5 | 0 | - | 0 | - | 28 |
| 6 | 2 | 0.74 | 10 | 0.94 | 386 |
| 7 | 11 | 0.96 | 20 | 0.79 | 509 |
| All | 47 | 0.82 | 71 | 0.86 | 2190 |

### Important substructure features

For those models where a small feature set was sufficient to reach ROC-AUC values above 0.8, the specific features were further analyzed. Each feature is described by an enrichment analysis (epitope coverage and fold-enrichment). For a detailed explanation of the metrics see Methods. The feature interpretation comprises the models for the T cell epitopes of the fatty acid derivatives (cluster 2), the T and B cell epitopes of the glucoside / oligosaccharide derivatives (cluster 4) and the T and B cell epitopes of the nucleobase-containing molecular entities (cluster 5).

#### Cluster 2 - fatty acid derivatives

The significant fingerprint features for T cell classification of fatty acid derivatives are listed in Table 4, with the corresponding substructures shown in Figure 8.

**Table 4.**
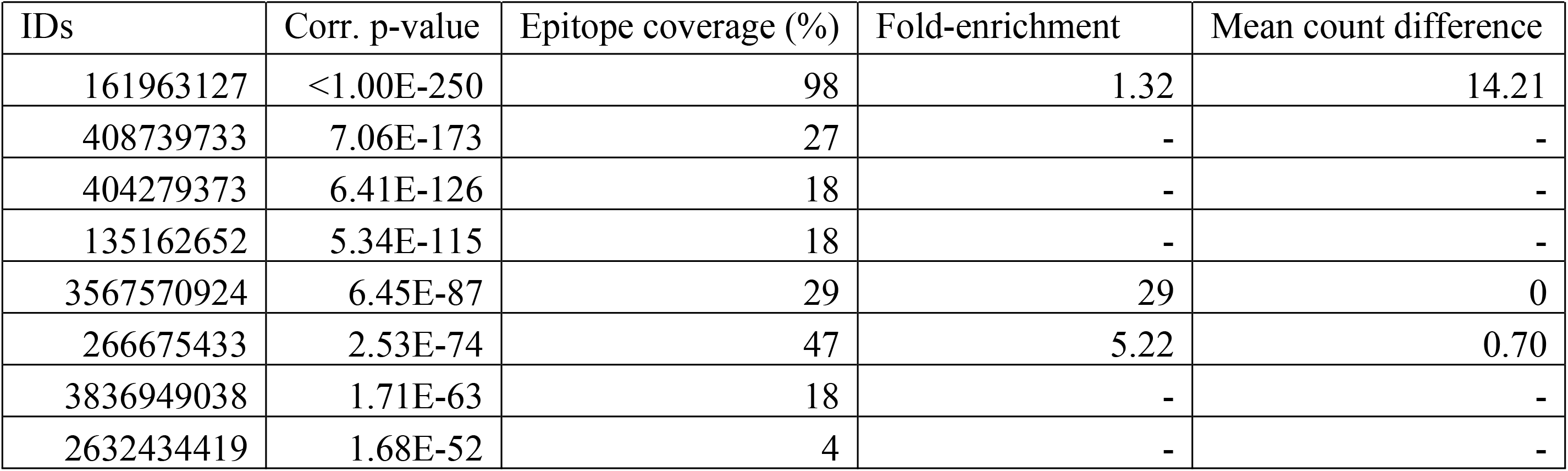
Most important fingerprint features for the prediction of T cell epitopes of the fatty acid derivatives (cluster 2). The fingerprint feature IDs correspond to Figure 8. The corr. p-value is based on the hypothesis (H_0_), that the feature count is equally distributed in the epitopes and the background. For explanation of other feature-specific metrics see Methods. For those features where no examples are present in the background dataset, the fold-enrichment and mean count difference cannot be computed.

**Figure 8.**
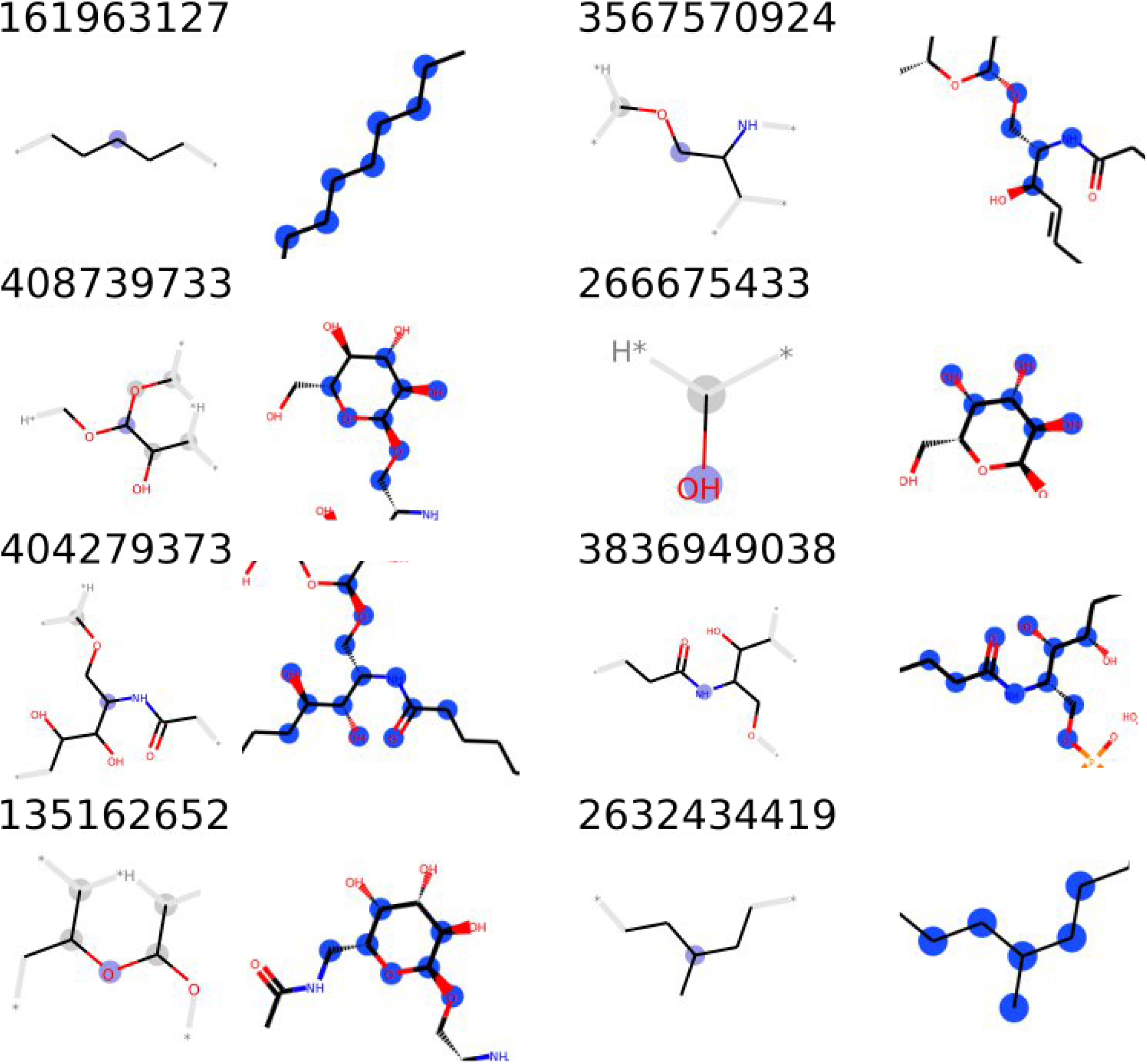
Substructures of most significant fingerprint features for the classification of T cell epitopes of the fatty acid derivatives (cluster 2). A depiction of each feature is shown on the left and a representation of the feature in an example molecule on the right. The statistics of the features are shown in Table 4.

The most important fingerprint feature (ID: 161963127) represents a carbon chain substructure. This substructure can be found in almost all molecules (epitopes and background) in this cluster (Fold-enrichment: 1.32). The significant difference is given by the mean substructure count of 14.21. Most epitopes possess much longer fatty acid chains than the ChEBI background molecules in the cluster.

Most of the other substructures can be associated with the attachment of a single sugar moiety to the fatty acid molecules. The most significant of those features (ID: 408739733) can be found in 27% of the epitopes, but not at all in the ChEBI background dataset.

#### Cluster 4 - glucoside / oligosaccharide derivatives

The T cell epitopes of the glucoside/oligosaccharide derivatives can be classified with high accuracy based on only one substructure (see Table 5). Surprisingly, this is the same fingerprint feature (ID: 161963127) as the key substructure for the T cell epitopes of the fatty acid derivatives.

**Table 5.**
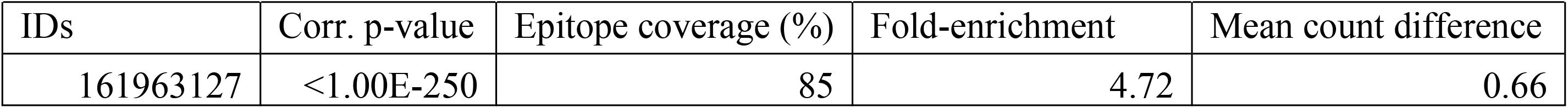
Most important fingerprint feature for the prediction of T cell epitopes of the glucoside/oligosaccharide derivatives (cluster 4). The fingerprint feature corresponds to Figure 9.

| IDs | Corr. p-value | Epitope coverage (%) | Fold-enrichment | Mean count difference |
| --- | --- | --- | --- | --- |
| 161963127 | <1.00E-250 | 85 | 4.72 | 0.66 |

A histogram of the substructure distribution in the epitopes and the ChEBI dataset is shown in Figure 9. The vast majority of known T cell epitopes (85%) have a long carbon chain attached to the glucoside, which can only be found in 18% of the overall ChEBI glucosides.

**Figure 9.**
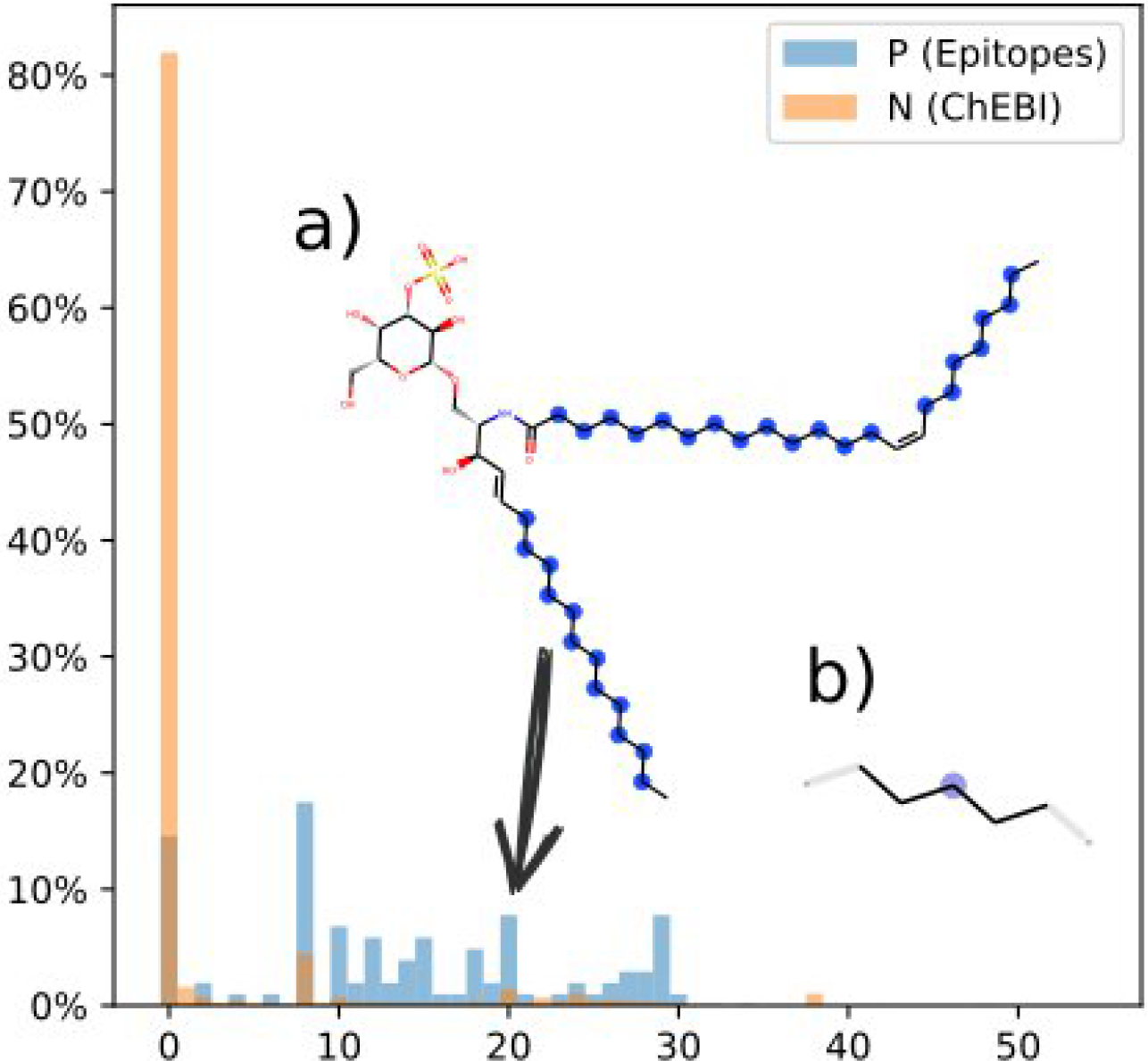
Histogram of the fingerprint feature (ID:16163127) count responsible for T cell prediction of the glucoside/oligosaccharide derivatives (cluster 4). The vast majority of epitopes have a long fatty acid chain attached to the glycoside. (a) Example molecule with 20 fingerprint features. (b) Depiction of the fingerprint feature.

The model for the B cell epitopes of glucoside/oligosaccharide derivatives requires 8 features to reach a ROC-AUC of 0.8 (see Figure 10). The statistics of these features are shown in Table 6.

**Table 6.** Most important fingerprint features for the prediction of B cell epitopes of the glucoside/oligosaccharide derivatives (cluster 4). The fingerprint feature IDs correspond to Figure 10.

| IDs | Corr. p-value | Epitope coverage (%) | Fold-enrichment | Mean count difference |
| --- | --- | --- | --- | --- |
| 161963127 | 1.17E-216 | 18 | 0.90 | -8.43 |
| 266675433 | 1.18E-188 | 100 | 1.00 | 2.79 |
| 2456262944 | 1.04E-137 | 94 | 1.00 | 1.71 |
| 951226070 | 2.11E-105 | 1 | 0.06 | 1.20 |
| 411967733 | 1.06E-86 | 64 | 1.78 | 0.13 |
| 784020300 | 4.72E-86 | 57 | 1.90 | 0.25 |
| 26234434 | 1.51E-77 | 0 | 0.00 | -0.95 |
| 1858577693 | 2.46E-74 | 35 | 1.94 | 0.36 |
| 161963127 | <1.00E-250 | 85 | 4.72 | 0.66 |

**Figure 10.**
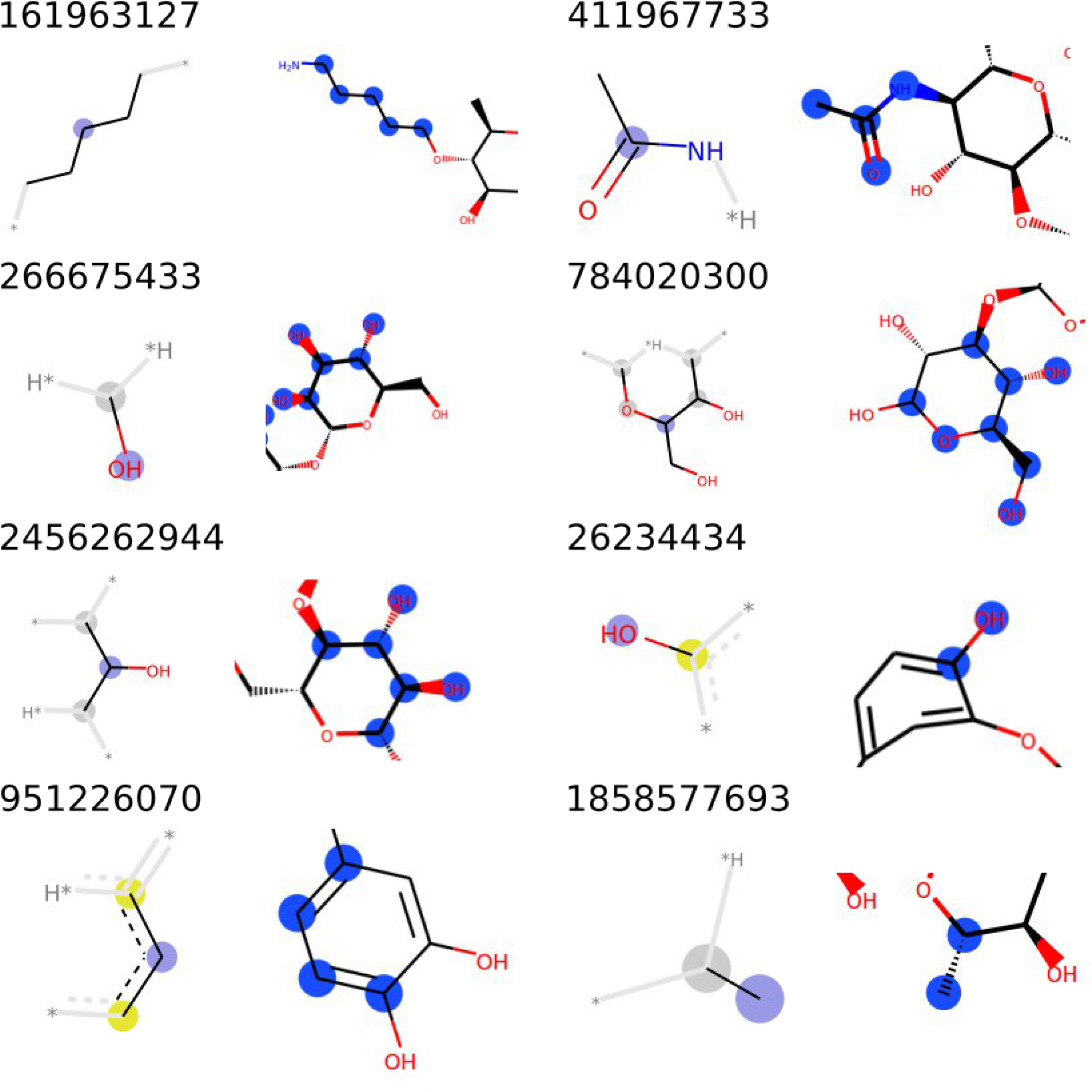
Substructures of most significant fingerprint features for the classification of B cell epitopes of the glucoside/oligosaccharide derivatives (cluster 4). A depiction of each feature is shown on the left and a representation of the feature in an example molecule on the right. The statistics of the features are shown in Table 6.

The most important substructure is again represented by the fingerprint feature with the ID 161963127. Nevertheless, the fatty acid attachment is much less common for the B cell epitopes (18%) as compared to the T cell epitopes (85%). Surprisingly, the B cell epitopes that possess this fatty acid attachment tend to have shorter chains as compared to the background molecules of the cluster (mean count difference: −8.43).

The other fingerprint features correspond either to specific sugar moieties (IDs: 26675433, 2456262944, 784020300) or aromatic substructures (IDs: 951226070, 26234434). While the sugar moieties can be found predominantly in the epitopes, the aromatic entities are negatively correlated to epitope activity. Another substructure that is enriched in the epitope dataset is given by the feature with the ID 411967733, a secondary amide.

#### Cluster 5 - nucleobase-containing molecular entities

Most features of the B cell epitopes of nucleobase-containing molecules can be associated with common nucleobases (see Figure 11). The classification decision can be explained by the substructure count difference, meaning that the epitopes tend to have more nucleobases than the background molecules of the cluster (see Table 7).

**Table 7.** Most important fingerprint features for the prediction of B cell epitopes of the nucleobase-containing molecular entities (cluster 5). The fingerprint feature IDs correspond to Figure 11.

| IDs | Corr. p-value | Epitope coverage (%) | Fold-enrichment | Mean count difference |
| --- | --- | --- | --- | --- |
| 151749292 | <1.00E-250 | 32 | 16.00 | 5.29 |
| 10565946 | <1.00E-250 | 80 | 1.23 | 8.77 |
| 1026924773 | <1.00E-250 | 7 | 3.50 | 18.96 |
| 77544489 | 1.73E-231 | 45 | 3.75 | 2.77 |
| 2245384272 | 7.80E-228 | 99 | 1.03 | 5.37 |
| 422715066 | 2.00E-218 | 36 | 2.40 | 4.30 |
| 5685888 | 1.39E-129 | 43 | 0.65 | 6.25 |
| 95089064 | 6.80E-125 | 31 | 3.10 | 2.67 |
| 151749292 | <1.00E-250 | 32 | 16.00 | 5.29 |

**Figure 11.**
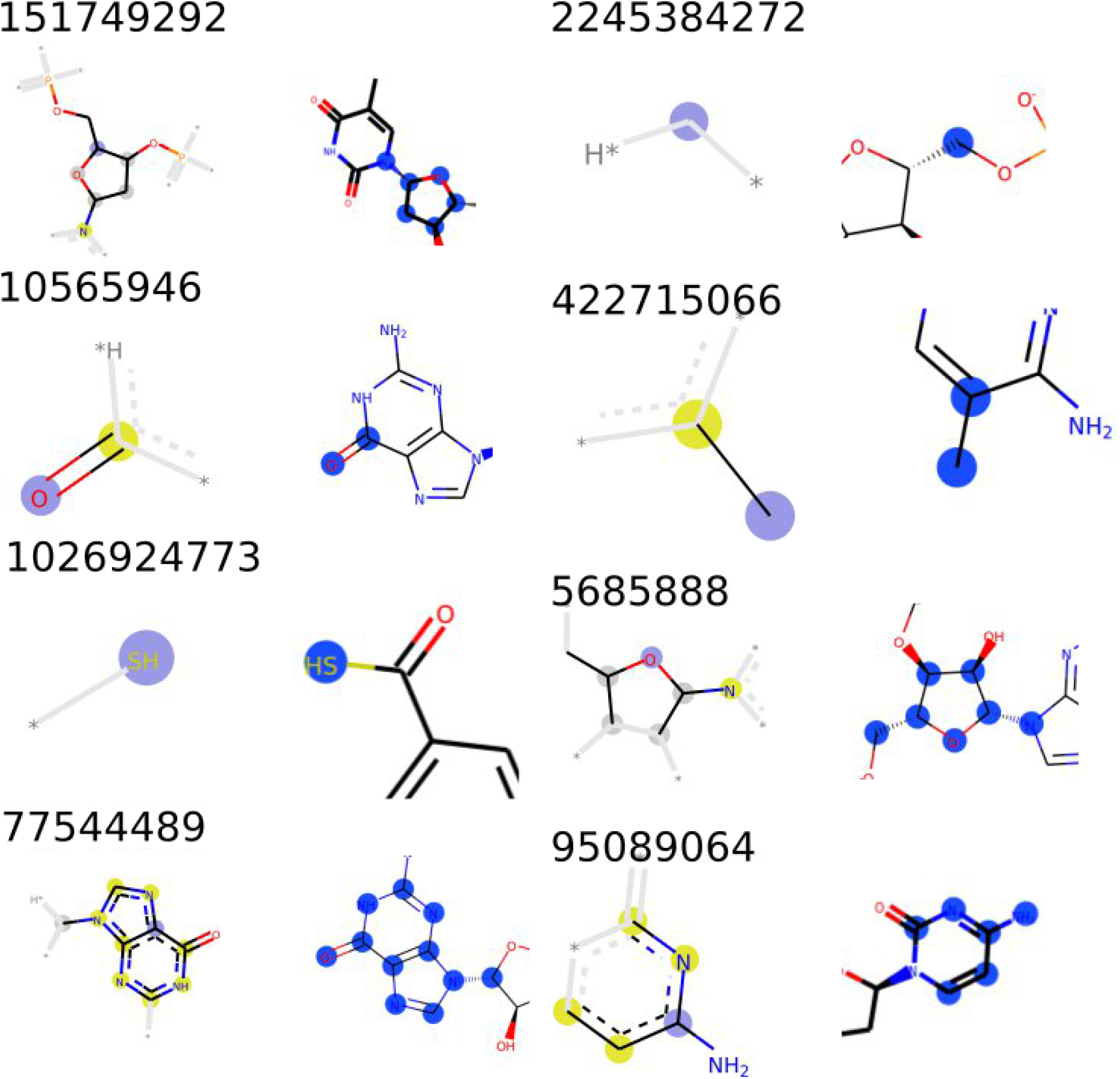
Substructures of most significant fingerprint features for the classification of B cell epitopes of the nucleobase-containing molecular entities (cluster 5). A depiction of each feature is shown on the left and a representation of the feature in an example molecule on the right. The statistics of the features are shown in Table 7.

The T cell epitopes can be classified solely on the presence of a single carbon chain moiety at the phosphorus backbone of the nucleobase (see Figure 12). This substructure can only be found in the epitope dataset and not in the background molecules (see Table 8).

**Table 8.** Most important fingerprint feature for the prediction of T cell epitopes of the nucleobase-containing molecular entities (cluster 5). The fingerprint feature corresponds to Figure 12.

| IDs | Corr. p-value | Epitope coverage (%) | Fold-enrichment | Mean count difference |
| --- | --- | --- | --- | --- |
| 3994088662 | 1.41E-192 | 60 | - | - |

**Figure 12.**
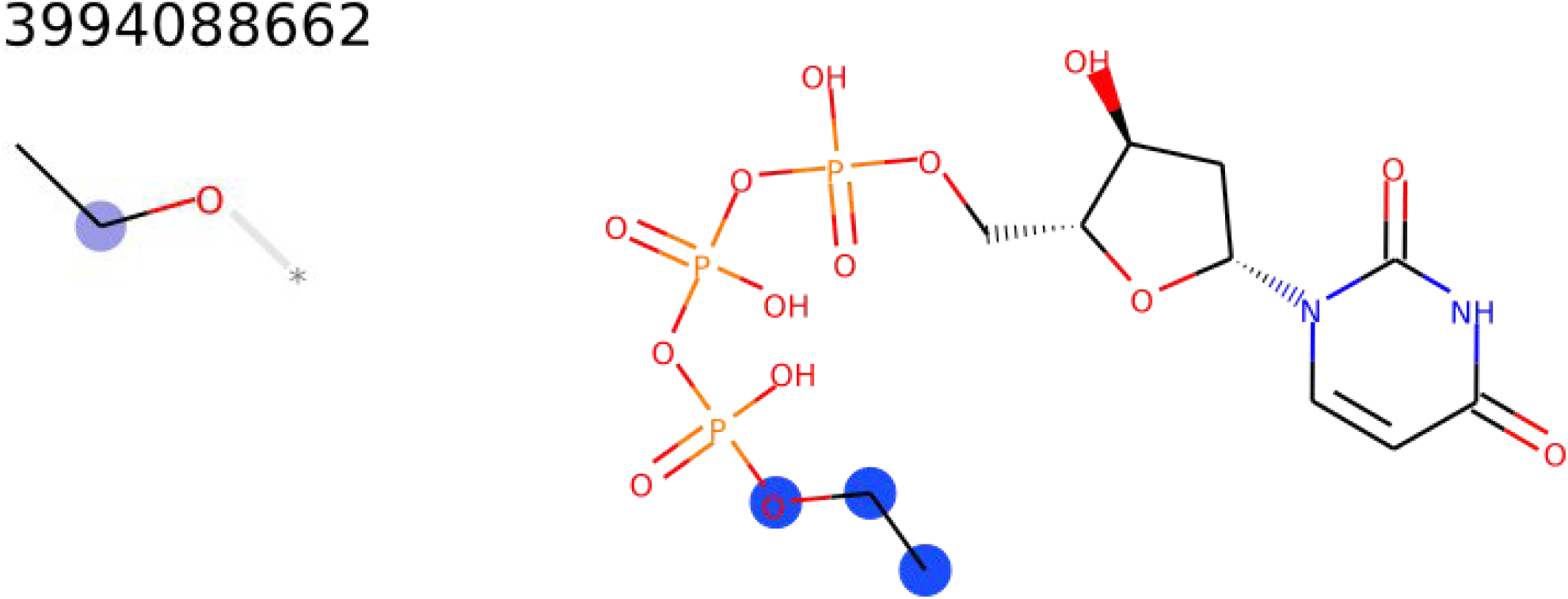
The feature responsible for the prediction of T cell recognition of the nucleobase-containing molecular entities (cluster 5). More than half of the known T cell epitopes of this cluster contain this feature, while no background molecule of the cluster has this moiety (see Table 8).

### Prediction tool

To allow the investigation of epitope activity of non-peptidic molecules, a prediction tool was developed (http://tools-staging.iedb.org/np_epitope_predictor). The tool takes a simplified molecular-input line-entry system (SMILES) representation of a molecule as input and performs a two-step analysis. First, the molecular class of the compound is predicted. In addition to the class membership, BiNChE statistics of the given class are shown (comp. Table 1). Second, the likelihood that a molecule could be an epitope binding to B cell or T cell receptors is predicted. The RF models built with the highest feature set were chosen for this task. For each epitope type, the significant fingerprint features found in the molecule are shown.

Furthermore, the fingerprint features of the 5 most similar epitopes from the IEDB are listed next to the query molecule, which allows for direct comparison of the substructures. The feature similarity is computed using Euclidean distance of the fingerprint feature counts. Moreover, the overall Tanimoto similarity, using unfolded Morgan count fingerprints (radius: 3, chiral) between the query molecule and the target epitopes, is computed. A link to the ChEBI entry of the query molecule is provided, which allows further investigation of similar epitopes. It is planned to update the tool regularly to allow for an analysis of non-peptidic epitopes based on the current state of the IEDB.

The tool was built as a small python application, that is controlled via a web interface based on the Django [25] web framework. The code of the application is open-source and available via (https://github.com/IEDB/NP_epitope_predictor) under the NPOSL-3.0 license.

## Discussion

### Clustering into homogeneous molecular subsets

To the best of our knowledge, this is the first attempt to use unfolded Morgan fingerprint features for the training of machine learning algorithms with a large dataset (> 40,000 molecules). The presented two-step approach could be used as a basis for the implementation of a general clustering-classification algorithm for molecular classification problems.

Most of the computed clusters contained uniform molecule sets, which could be described with BiNCHE ontology analysis. Nevertheless, the entire ChEBI database contains a diverse range of small biologically relevant molecules and, subsequently, some molecules are difficult to aggregate. This could be observed especially for cluster 6, which is a collection of diverse small molecules, simply because molecules with few substructure features are aggregated by k-means clustering logic. Interestingly, the most distinct cluster, cluster 3, which comprises exclusively CoA derivatives, does not contain any known epitopes. It can be hypothesized that the lack of CoA-related epitopes is due to the involvement of CoA in various crucial biological functions, such as fatty acid synthesis and the citric acid cycle. An immune response against such a biologically vital molecule might be generally suppressed due to negative selection during immune cell maturation. The putative suppression of immune cell responses against vital self-molecules should be further investigated, for example based on co-factor-related compounds.

### Epitope prediction

The radius parameter of the Morgan fingerprint algorithm led to different classification performances concerning the observed clusters. We settled for a radius of 3 as a compromise for the subsequent analysis, since most models showed good performance with this option. We decided that the additional inclusion of different radius options for the model comparison would have made the subsequent investigation too complex. Nevertheless, the strong performance fluctuation of clusters 1 and 2 concerning the radius option may be related to fatty acid and glycolipid molecules, which possess repetitive molecular characteristics.

Different machine learning models were compared for the ability to predict B cell- or T cell-related epitope activity. The RF and NN models could outperform the k-NN models in all categories. The hyperparameters of the models were not optimized, because this could have led to overfitting of the models to the training samples. The overall experimental set-up was tested using the dummy RF classifiers (trained on randomly assigned positive samples). These classifiers showed the expected ROC-AUC of 0.5 for all instances, confirming that no information leak or stratification bias occurred with the given dataset.

The performance of all models increased with the number of features used to train the models. In most cases, the ROC-AUC approached a plateau, indicating that more features did not lead to further information gain. The increase of features for a fixed number of training samples can often lead to a performance drop, due to the addition of potentially uninformative features - referred to as the Hughes phenomenon [26]. This drop could only be observed in very few cases, such as the RF model for cluster 1 (both epitope types). The lack of a performance drop could be explained by the nature of the Morgan fingerprint features. Given that the Morgan fingerprints are based on correlating substructures; additional features are likely to be redundant but not uninformative. This would explain the plateau for high feature numbers.

The RF classifiers were compared against Tanimoto-similarity based classifiers. The Tanimoto-similarity based classifiers can be regarded as reference models to estimate the performance of memorizing the overall molecular structure as opposed to generalization. Indeed, it can be observed that for most clusters the similarity models and the RF models perform comparably, given enough features. Only the RF models built for the B cell epitopes achieved significantly better ROC-AUC scores in some cases. This could be explained by the higher number of positive training samples for the B cell epitopes.

Because the epitopes were manually collected, it is conceivable that a certain amount of sampling bias can be attributed to the dataset. Overfitting to similar molecules is a frequently encountered problem in molecular encoding based machine learning tasks [27]. A straightforward approach to avoiding overfitting is the choice of models trained with few features regarding the training samples. Consequently, the models that yielded good performance even for low feature sets are most likely to allow for correct predictions of novel molecules. This was achieved for clusters 4 and 5 and the T cell epitopes of cluster 2. Those models are most likely to allow for predictions of epitopes based on specific features instead of overall molecular similarity.

A common approach to estimate the performance of a classifier on novel samples is the usage of an independent test dataset, such as one that is not used in the model building process. Therefore, the updated samples from the IEDB and ChEBI, which were collected after the initial training of the classification tool, were chosen. Although some clusters had no representative positive samples in the test dataset, the overall performance allowed for the classification of B and T cell epitopes with high ROC-AUC scores.

### Interpretation of substructure features

The most important factor to classify fatty acid T cell epitopes (cluster 2) can be associated with the length of the fatty acid. Another important feature is given by the attachment of a sugar to the fatty acid chain. This finding is consistent with the features observed for the glucoside/oligosaccharide derivatives (cluster 4). The attachment of long carbon chains to the glycoside is highly correlated to T cell activation. In summary, it can be concluded that long fatty acid chains, especially with specific saccharide moieties, are the most significant indicator for T cell recognition. The T cell recognition of glycolipids by CD1 proteins has been described by Young *et al.* [28]. It was shown that T cells can specifically discriminate various moieties attached to the fatty acid. This is consistent with the findings derived from the observed models.

The most important feature for glucoside/oligosaccharide derivative (cluster 4) B cell epitope classification is also given by the chain length of fatty acid attachments. Surprisingly, the chain length is negatively correlated with B cell activation (observed in 18% of the epitopes). This means that glycoside molecules, which do have a short fatty acid, tend to be recognized by B cells, while longer fatty acid attachments are not. In general, the B cell epitope classification of cluster 4 is rather difficult to interpret, because various sugar and aromatic substructures are involved. The most intuitive finding is given by the feature with the ID 411967733 (see Figure 10). The corresponding substructure represents a secondary amide which is present in all samples of cluster 4 (Fold-enrichment: 1). But the epitopes have 2.79 times more instances of this moiety. Secondary amides are part of the building blocks (N-acetylglucosamine and N-acetylmuramic acid) of peptidoglycans as well as lipopolysaccharides (LPS). It is not surprising to find such a feature enriched in the epitope dataset, because these moieties (which are found in the cell walls of bacteria) are common non-self-microbial signatures [29] found in pathogen-associated molecular patterns (PAMPs). Although PAMPs are often associated with the initial defense provided by the innate immune system [30], they are also commonly encountered as bacterial-specific antibody counterparts [31–33].

The B cell epitopes of the nucleobase-containing molecules (cluster 5) can be classified based on the number of nucleobases. This finding may be attributed to the data collection process of the ChEBI and IEDB databases. ChEBI does not curate nucleobases derived from normal metabolism (e.g., DNA and RNA fragments), while the IEDB includes any nucleobase-containing entity with a positive immune cell assay. This could have led to an accumulation of molecules with longer nucleobase chains in the IEDB dataset.

The T cell epitopes of the nucleobase-containing molecules (cluster 5) could be classified based on only one substructure feature. This substructure, an ethyl ester attached to the phosphor part (ID: 399408862), could be found in 60% of the epitopes and none of the 187 background molecules. An investigation of the samples revealed that all the epitopes were collected from the same study by Tanaka *et al.* [34]. On the one hand, this finding highlights the potential risk of data collection bias for machine learning models built from small datasets; therefore, we evaluated the final models using a test dataset, where the retrieval of the samples from different studies was ensured. On the other hand, the finding supports the power of the developed method because the study by Tanaka *et al.* showed that monoethyl phosphates mimic mycobacterial antigens. Our model could derive the importance of this moiety based on the provided samples.

## Conclusions

In the presented work, the first general attempt was made to predict the recognition of non-peptidic molecules by B cell and/or T cell receptors. The generated models, as implemented in the web server, allow for a comprehensive analysis of non-peptidic molecules regarding epitope activity - despite the limitations of the available training dataset. The implemented prediction, as well as the shown similarity to known epitopes, allows users to judge whether the prediction is based on specific molecular features or on overall molecular similarity. The noteworthy ROC-AUC scores for the independent test dataset demonstrate the general usability of the software to investigate the epitope activity of novel non-peptidic molecules. The provided framework allows for a continuous update of the generated models and calculated decision rules with each major update of the IEDB. Thus, our framework provides a solid basis for the community to further explore non-peptidic epitopes.

## Methods

### Dataset

An unambiguous negative dataset of non-peptidic epitopes is problematic to collect because immunogenic reactions can be heterogeneous and limited to individuals; furthermore, a multitude of molecules was not systematically analyzed for immunogenic effects, yet. Therefore, the entirety of molecules in the ChEBI database [22] was assigned as a background dataset (downloaded: May 11, 2020). ChEBI has both manually curated and automatically assigned molecular structures. Only the molecules curated by the ChEBI team (marked with three stars in the database) were used. Positive structures tested in B cell and/or T cell assays were downloaded from the IEDB via a web-interface query (https://www.iedb.org/; downloaded: May 11, 2020). All structures were parsed using the cheminformatics python package RDKit [35], and those with duplicate SMILES [36] were removed. The final dataset included 42,643 ChEBI background molecules; 579 molecules tested positive in T cell assays and 2,140 molecules tested positive in B cell assays.

Molecules that were added to the IEDB or ChEBI databases after the May 11, 2020 were used as an independent test dataset to benchmark the developed prediction tool using samples that were not used in the cross-validation. The test dataset included 2,190 ChEBI background molecules; 71 molecules tested positive in T cell assays and 47 molecules tested positive in B cell assays.

### Molecular fingerprints encoding

The molecules were encoded into vectors by applying the Morgan fingerprint algorithm, also referred to as Extended-connectivity fingerprint (ECFPs) [23], using RDKit. The Morgan algorithm creates substructures of molecules by generating circular patterns with a certain radius from each atom in the molecule. Resulting substructures are used to set bit features (1 if the substructure is present in the molecule and 0 if the substructure is absent) or count features (the number of occurrences of the substructures in the molecule) in an array referred to as the fingerprint of the molecule.

The fingerprint can be folded, leading to a reduction of the data size, which is useful when the sample size is large since it requires less computing power [37]. However, the folding of the fingerprint can lead to bit collision (multiple substructures set the same fingerprint features), which does not allow for an interpretation of the substructures in folded fingerprints.

The RDKit implementation of the Morgan fingerprint allows for the creation of different features considering molecular chirality. The radius and chirality options were benchmarked using RF classifiers as described in the epitope prediction part of the Methods.

### Clustering into homogeneous molecular subsets

The ChEBI database represents a large collection of diverse molecules. Molecular subgroups, such as fatty acids, carbohydrates, and small molecules are likely to activate B cells and T cells with different mechanisms. For example, whereas lipids and glycolipids are known to be presented by MHC-like CD1 molecules [10,11], carbohydrates can be recognized by anti-carbohydrate antibodies [38]. Furthermore, haptenic determinants rely on the mediation of carrier proteins for immune cell activation. To account for different immunogenic mechanisms, it is required to investigate the molecular groups independently. Since clear distinction features cannot be derived for such a finely differentiated dataset, we decided to separate the molecular groups using k-means clustering [39].

From a technical point of view, the clustering into more homogenous molecular subsets was also advantageous. To avoid bit collision, we wanted to use unfolded Morgen fingerprints for the classification models. However, the entire ChEBI dataset would generate a large amount of fingerprint features, exceeding the computation power of conventional computers. In contrast, the molecular subsets generated far fewer features and could be used for prediction and statistical interpretation.

The ChEBI dataset was converted into folded Morgan bit fingerprints (1024 bits, radius: 3, non-chiral). The molecules were clustered using the k-means clustering algorithm implemented in the machine learning python package Scikit-learn [40]. The optimal number of clusters was determined using the elbow method [41]. The clusters were described using BiNChE ontology enrichment analysis [24], allowing for the interpretation of the clusters based on functional compound classes. BiNChE returns a table with corrected p-values, the fold-enrichment (ratio between the enrichment in the selected samples and enrichment in the background samples), and the sample coverage (percentage of the molecules that contain the ontology term) of significant ontology terms.

To visualize the clusters in a 2D representation, the vectors were transformed into 2 orthogonal components that explain the maximum amount of variance using Principal Component Analysis (PCA) [42] as implemented in Scikit-learn.

### Epitope prediction

Unfolded Morgan count fingerprints (radius: 3, chiral) were used to train the classification models. We considered count features as advantageous to bit features since there are various examples where repetitive molecular structures (e.g., fatty acids) play an important role in immune cell recognition [28]. These molecules would lead to identical bit-based fingerprints, but different count-based fingerprints.

The fingerprints were computed for each cluster separately. Molecules with identical fingerprints were removed from the dataset. The fingerprints were trimmed to include only those features which occurred in at least 10 molecules. Specific count features in a Morgan fingerprint can highly correlate. To derive clear decision rules, correlating fingerprint features, which exceeded a Pearson Correlation Coefficient (PCC) of more than 0.8 to any other feature, were removed.

For each cluster, two machine learning models were compiled that predict the probability of a molecule to act as a B cell or T cell activating epitope. Different models were created and compared against each other. RF, k-NN, and NN algorithms with default parameters as implemented in Scikit-learn were used. For the RF models, 100 iterations were selected. Furthermore, dummy RF models were designed to validate the experimental set-up. For the dummy models, the positive samples were assigned from the background by random shuffling. The percentage of positive samples was identical to the real epitope percentage in each cluster. The dummy classifier should demonstrate that no learning process can be achieved from arbitrary samples. To analyze the feature importance, different fingerprint feature sets were selected using the chi-squared feature selection approach implemented in the Scikit-learn “SelectKBest” algorithm. The algorithm is described in the article by Cressie *et al.* [43]. All classifiers were benchmarked using a repeated (3 times) 5-fold stratified cross-validation. The classifier performance was compared using the ROC-AUC metric.

### Benchmark against Tanimoto similarity-based classifiers

The RF classifiers were compared against a classifier that uses a Tanimoto similarity-based prediction approach. The similarity classifiers were designed as follows: for each molecule, the Tanimoto similarity to all known epitopes in a cluster was calculated using unfolded Morgan count fingerprints (radius: 3, chiral). The highest similarity was then assigned as a score of this structure to be an epitope.

All similarity-based classifiers were benchmarked using repeated (3 times) 5-fold stratified cross-validation. The classifier performance was estimated by computing the ROC-AUC metric.

### Fingerprint substructures

To extract fingerprint features that are important to distinguish epitopes from the background, a statistical investigation of the fingerprint distribution for each cluster was carried out. The investigation was focused on those fingerprints, for which classification models with high performance (> 0.8 ROC-AUC) could be built using a set of no more than 8 features.

From the chi-squared feature selection approach, the p-values were extracted. For each feature the null hypothesis was tested to ensure that the feature count of the epitopes was selected from the same distribution as for all ChEBI structures without epitopes. The p-value can be used as a measure for the feature importance. The p-values were additional Bonferroni corrected to account for the multiple comparison problem [44].

Additionally, feature-specific statistics were computed, which allowed for further interpretation of the features. The statistics for each substructure include the epitope coverage, the fold-enrichment of the feature in the epitopes, and the mean count difference. The epitope coverage shows the percentage of the epitopes that have the substructure. The fold-enrichment is computed by dividing the sample percentage by the percentage of background molecules that have the substructure. The mean count difference is computed by the difference between the mean of the substructure count for the epitopes and the background. Molecules that did not have a specific substructure were not used for the computation of the mean count difference.

Figures of the substructures corresponding to the fingerprint features were computed using the custom RDKit function. Thereby the fingerprint gets extracted based on the occurrence in a certain example molecule.

## Notes

### Competing Interest Statement

The authors have declared no competing interest.

